# Electrospun Cellulose-Silk Composite Nanofibres Direct Mesenchymal Stem Cell Chondrogenesis in the Absence of Biological Stimulation

**DOI:** 10.1101/434316

**Authors:** Runa Begum, Adam W. Perriman, Bo Su, Fabrizio Scarpa, Wael Kafienah

## Abstract

Smart biomaterials with an inherent stimulating capacity that elicit specific behaviours *in lieu* of biological prompts would prove advantageous for regenerative medicine applications. Specific blends of the natural polymers cellulose and silk cast as films can drive the chondrogenic differentiation of human bone marrow mesenchymal stem cells (hMSCs) upon *in vitro* culture. However, the true potential of such biomaterials for cartilage tissue engineering can be realised upon its three-dimensional fabrication. In this work we employ an electrospinning technique to model the *in vivo* nanofibrous extracellular matrix (ECM). Cellulose and silk polymers at a mass ratio of 75:25 were regenerated using a trifluoroacetic acid and acetic acid cosolvent system. This natural polymer composite was directly electrospun for the first time, into nanofibers without post-spun treatment. The presence and size of fibre beading was influenced by environmental humidity. The regenerated composite retained the key chemical functionalities of its respective components. Biocompatibility of the natural polymer composite with hMSCs was demonstrated and its inherent capacity to direct chondrogenic stem cell differentiation, in the absence of stimulating growth factors, was confirmed. This physical chondrogenic stimulation was countered biochemically using fibroblast growth factor-2 (FGF-2), a growth factor used to enhance the proliferation of hMSCs. The newly fabricated scaffold provides the foundation for designing a robust, self-inductive, and cost-effective biomimetic biomaterial for cartilage tissue engineering.

## 1. Introduction

Biomaterials used in tissue engineering act as a scaffold for cell growth *in vitro* and cell delivery *in vivo*, supporting the cells whilst they develop and establish their own *de novo* scaffold – the extracellular matrix (ECM). The *in vivo* ECM largely consists of high-strength fibrous collagens embedded in a hydrated proteoglycan matrix, allowing cell to cell communication and directed tissue formation. The collagen fibres, composed of nanometre-scale multifibrils, form 3D macroscopic tissue architectures, which vary between tissue types, with fibre diameters ranging from 50-500 nanometres [1]–[3]. Biomaterial fabrication processes have begun focusing on mimicking this nanoscale morphology, and electrospinning has been widely utilised for the development of 3D fibrous scaffolds, particularly in the field of cartilage tissue engineering [4]–[10].

Electrospinning involves the fabrication of polymer fibres through the exploitation of electrostatic forces. Both natural and synthetic polymer sources have been employed in such technology, producing fibres that range in diameters from a few nanometres to several micrometres [11]. Electrospinning, therefore, allows the assembly of an artificial ECM retaining the cells native structural milieu. Compared to other fibre spinning processes, electrospinning permits the generation of long fibres with smaller diameters and higher surface area-to-volume ratios. It is not surprising therefore, that such a method has been widely explored for various applications [11]. In the context of tissue engineering, fibrous materials would be advantageous to resident cells, by supporting the efficient exchange of nutrients, gases and waste products. Trifluoroacetic acid (TFA) was identified as a common solvent for use in electrospinning, having previously been used to electrospin silk fibroin [12], cellulose [13], and recently, blends of these polymers. Guzman-Puyol *et al.* directly electrospun blends of microcrystalline cellulose and silk fibroin in a 50:50 ratio [14]. They were unable to fabricate blends with a higher cellulose content. Furthermore, acetic acid (AcOH) can be incorporated as a cosolvent with TFA to help reduce solution viscosity and enhance solution spinnability and has been used for the electrospinning of cellulose-chitosan composite fibres [15] and as an aqueous solvent for electrospinning cellulose acetate [16] and chitosan nanofibres [17]. It has also been used to control the viscosity of solutions in TFA when casting neat cellulose films [18] and blend films of cellulose and chitosan [19], [20].

Silk fibroin and cellulose serve a common purpose in their respective native environments as structural protein-based support [21], [22]. These highly abundant, natural, renewable polymers make ideal candidates for use in medical applications, not least due to their biocompatibility. Their chemical structures ensure a high level of mechanical integrity making them some of nature’s most attractive polymers. *B.mori* silk fibroin protein consists of anti-parallel ßsheets stacked together by strong intra- and intermolecular hydrogen bonding interactions. Similarly, cellulose comprises a polysaccharide chain of ß1-4 glycosidic bonded anhydroglucose units with intramolecular hydrogen bonds between the repeating units and intermolecular hydrogen bonding between polysaccharide chains, resulting in a highly ordered linear, rigid structure [21]. Silk fibroin has been electrospun in a wide range of solvent systems [23], [24]. Conversely, cellulose has only been directly electrospun in TFA [13]. Its derivatives, particularly cellulose acetate, have been widely electrospun, but require post-spinning treatments to remove the acetyl group and regenerate pure cellulose fibres [25]. The requirement of post-spinning treatments following electrospinning may not be feasible when fabricating cellulose-silk fibrous blends.

Whilst much research has investigated electrospun fibrous mats for cartilage tissue engineering, a limited number of researchers have used natural polymers and all require the presence of stimulating factors to direct cell behaviour [4], [5], [7]–[10], [26]. We have demonstrated recently that a specific blend of cellulose and silk was able to drive the chondrogenic differentiation of human mesenchymal stem cells (hMSCs) [27]. Human MSCs seeded onto cast films of blended cellulose and silk in a 75:25 mass ratio, in the absence of stimulating factors, expressed chondrogenic genes and deposited cartilaginous ECM proteins. This specific blend therefore shows promise for cartilage tissue engineering in the treatment of degenerative joint diseases. The blends true potential for clinical application can only be realised once this specific 2D composite is fabricated into a 3D biomimetic configuration.

Herein we describe, for the first time, the direct electrospinning of cellulose-silk in a 75:25 mass ratio using a TFA-AcOH cosolvent system. We demonstrate the biocompatibility of these novel composite nanofibres with hMSCs. Not only did the material support the growth of hMSCs, they also directed their differentiation toward the chondrogenic lineage, in the absence of stimulating factors. The chondrogenic stimulation could be countered biochemically by increasing the concentration of fibroblast growth factor-2 (FGF-2); a growth factor commonly used to enhance the proliferation of hMSCs [28], [29] [30].

## 2. Materials and Methods

### 2.1. Fabrication of Electrospun Composite Fibres

*Bombyx mori* silk (Aurora Silk, Portland, USA) was degummed to obtain silk fibroin fibres as follows; fibres were cut into 1 cm pieces and boiled in 0.02 M Na_2_CO_3_ solution. Fibres were then washed thoroughly with distilled water to ensure removal of sericin proteins and left to dry overnight in a fume hood. Use of silk fibroin shall be referred to as silk from herein. Cellulose from wood pulp, degree of polymerisation (DP) 890, was purchased as sheets (Rayonier Inc., Jacksonville, USA) and ground to a powder. Trifluoroacetic acid (TFA, 99 %) and glacial acetic acid (AcOH, ≥99.85 %) were purchased from Sigma Aldrich (Dorset, England).

A 4 wt% concentration blend solution of cellulose:silk in a 75:25 ratio was prepared in TFA. The polymers were weighed out into a glass vial containing a stir bar. Following solvent addition, vials were sealed and placed on a magnetic stirrer at steady speed and room temperature for 8 days. AcOH was added to the solution at 20 % (v/v), taking the initial volume of the solution (in TFA) as 80 %. The solution was thoroughly mixed and electrospun immediately. A horizontal electrospinning setup was used under a fume hood. The electrospinning apparatus was set-up as follows; the polymer solution was loaded into a syringe (5 ml Leur-Lok™, Beckton Dickinson, NJ, USA), and a stainless steel blunt-end 22 gauge needle attached (Precision glide, Beckton Dickinson, NJ, USA). This was then placed onto a syringe pump (PHD 2000 Infusion Pump, Harvard Apparatus, MA, USA), facing a flat metal plate covered with aluminium foil. A high voltage power supply (EL Series 1-30 kV, Glassman High Voltage Inc, EU) was connected to the needle tip and grounded at the metal foil (collector). The distance between the needle tip and the collector was kept at 10 cm. A 4 wt% concentration solution electrospun at a 1.0 ml/hr flow rate and 2.0 kV/cm voltage was identified as ideal; enabling a continuous electrohydrodynamic jet to be spun. All experiments were performed at room temperature. Although environmental temperature and relative humidity (RH) were not controlled, they were monitored. Values for each parameter were recorded at the beginning of electrospinning, then every ten minutes thereafter. Recordings were made over the electrospinning period using a hygrometer (Testo 608-H2 Humidity, Temperature & Dewpoint Hygrometer 120603).

### 2.2. Characterisation of Composite Fibres

Fibre morphology and diameter was assessed using a combination of scanning electron microscopy (SEM) and ImageJ software [31]. Fibrous mats were placed on a carbon pad and sputter coated with gold in argon, with a plasma current of 18 mA for 15 seconds, (SC7620 Mini Sputter Coater, Quorum Technologies, Kent, UK). The resulting coat thickness was 45.3 Å (4.53 nm). Samples were then imaged using a field emission gun scanning electron microscope (Zeiss EVO Series SEM, Carl Zeiss, UK) at 15 kV accelerating voltage and 24 mm working distance. SEM micrographs were loaded into ImageJ and fibre diameter/beading measured (ImageJ 1.x) [31]. The micrograph scale bar was used to set the image resolution in pixels/μm. Measurements were then performed adopting a two-point measurement approach. For each polymer type, at each electrospinning parameter, 10 SEM micrographs were taken. Fibre diameters were recorded by measuring distance across the width of fibres. Ten fibres were measured from each image, totalling 100 measured fibres per fibre type. Fibre beading was characterised in two ways; bead size and bead frequency. Bead size was characterised by measuring length (along the axis of the fibre) and width (across centre of bead). Five beads were measured, length and width of each, from four different micrographs, totalling 20 measured beads per fibre type. Bead frequency was measured by isolating a 100 μm^2^ area (10 μm × 10 μm) of the micrograph and counting the number of beads enclosed. This was done on a total of four separate areas taken from four different micrographs.

Chemical characterisation was performed on the nanofibrous mats using Fourier transform infrared spectroscopy (Spectrum 100 FTIR spectrometer, PerkinElmer, MA, USA). Analysis was performed in transmission mode across a spectral range of 4000-600 cm^−1^. Fibrous mats were analysed as well as the raw polymers to investigate any difference in chemistry following dissolution in the solvents and electrospinning. Spectra were generated three times per sample to ensure chemical homogeneity throughout samples.

### 2.3. Human Mesenchymal Stem Cell Culture

Mesenchymal stem cells (MSCs) were isolated from human bone marrow plugs recovered from patients undergoing complete hip replacement arthroplasty. Sample collection was carried out following local ethical guidelines in Southmead Hospital, North Bristol Trust. MSCs from five patients were used (*n*=5) in the study. Following isolation, MSCs were expanded in stem cell expansion medium consisting of low glucose Dulbecco′s modified Eagles medium (DMEM), 10 % (v/v) fetal bovine serum (FBS, Thermo Scientific Hyclone, UK), 1 % (v/v) Glutamax (Sigma, UK), and 10 % (v/v) penicillin (100 units/mL)/streptomycin (100 mg/ml) antibiotic mixture (P/S, Sigma, UK). The stem cell expansion medium was supplemented with 5 or 10 ng/ml fibroblast growth factor 2 (FGF- 2, PeproTech, UK). FGF-2 is known to enhance bone marrow MSC proliferation and chondrogenic differentiation potential [28], [29]. Cells were cultured at a density of 2.0×10^5^ cells per cm^2^ and incubated at 37 °C in a humidified atmosphere of 5 % CO_2_ and 95 % air. Media was changed every 2-3 days and cells were passaged upon reaching 80-90 % confluency. Cells used for all experiments were between passage 0 and 2.

### 2.4. Cell Loading on Composite Nanofibres

Composite nanofibrous mats were cut into 8 millimetre discs using a biopsy punch (Stiefel, Schuco Intl). Discs were disinfected as follows; aqueous 70 % ethanol (v/v) was added to tissue culture plastic containing nanofibrous discs for 30 minutes and then washed three times with sterile phosphate buffered saline (PBS) to remove any residual alcohol.

For fibronectin coated scaffolds; following the wash step, PBS was removed and 100 μg/ml human plasma fibronectin in PBS was added (Sigma, UK). Adsorption of human plasma fibronectin on polymer surfaces enhances cell adhesion [32]. For non-fibronectin coated scaffolds; following the wash step, PBS was removed, and scaffolds immersed in further PBS. Fibronectin and non-fibronectin coated scaffolds were then incubated at 37 °C in a humidified atmosphere of 5 % CO_2_ and 95 % air overnight. Following overnight incubation, scaffolds were transferred to 24-well ultra-low attachment plates (Corning, Costar, Sigma UK) containing sterile PBS. The PBS was then removed, enabling complete placement of nanofibrous mats at the bottom of the well. The ultra-low attachment plate wells are coated with an inert hydrogel layer that inhibits cell adhesion (Corning, Costar, Sigma UK). This would ensure cells added to the well would only interact with the scaffold and not bind to the well surface. Plates were covered and left at the back of the tissue culture hood to allow the scaffolds to dry (~2 hours). Stem cell expansion media (as described above) was then added to each well and plates transferred to the incubator (37 °C, 5 % CO_2_, 95 % air) until MSCs were ready for loading. Cells were plated at a density of 28×10^3^ cells per cm^2^ on scaffolds and tissue culture plastic controls. For cell loading on nanofibrous mats, MSC cell suspension was pipetted directly on top of the scaffold. Plates were then returned to the incubator and media changed every 2-3 days.

### 2.5. Characterising MSC Behaviour on Composite Nanofibres

Mesenchymal stem cell morphology was visualised on the nanofibrous composites using SEM. Cell-loaded materials were fixed in 4% paraformaldehyde then dehydrated in a series of ethanol solutions of increasing concentration. The samples were then dried using a critical point dryer (Leica EM CPD300) and prepared for SEM as discussed above.

MSC viability and proliferation on nanofibrous mats was assessed quantitatively using Alamar Blue (AB) assay (ThermoFisher, UK). The AB dye is a non-toxic fluorescent dye that undergoes a REDOX reduction in the presence of cellular growth, fluorescing and changing colour [33]. In the presence of metabolic activity, the dye undergoes a chemical reduction, changing from its non-fluorescent blue form (oxidised) to a fluorescent red form (reduced). The degree of this often-visible colour change (i.e. reduction) is indicative of the level of cell proliferation which is quantified using a spectrophotometer at 560 nm and 600 nm wavelengths (GloMax-Multi+ Microplate Multimode Reader with Instinct, Promega).

Nanofibrous mats (with and without fibronectin coating) were cultured with MSCs in stem cell medium supplemented with 5 ng/ml FGF-2. MSCs were also grown on standard 24-well tissue culture plastic (Sigma, UK) at the same cell density (28×10^3^ cells per cm^2^) and same media conditions, as a control. The AB assay was performed at days 1, 3 and 7 following initial cell loading (day 0) on the same wells. The percentage reduction of Alamar Blue was calculated following a standard protocol (alamarBlue^®^ protocol, Bio-Rad antibody). Following the Alamar Blue assay at day 7, MSC adhesion and viability was qualitatively assessed using the Live/Dead Viability/Cytotoxicity assay kit (Invitrogen, UK), following the manufacturer’s protocol. This assay is based on the principle that viable cells have intracellular esterase activity and plasma membrane integrity. Calcein AM is used to measure the former parameter. This non-fluorescent cell-permeant dye is converted to the intensely fluorescent Calcein (green) in the presence of esterase. Plasma membrane integrity is detected using Ethidium homodimer-1 (EthD-1), which can enter cells with damaged membranes and bind to nucleic acids, creating red fluorescence. EthD-1 is unable to enter the intact membranes of live cells. Cells were incubated for 30 minutes at room temperature with 2 μM Calcein AM (green fluorescent dye, live cells, 488 nm emission) and 4 μM Ethidium homodimer-1 (EthD-1, red fluorescent dye, 568 nm emission). Cells were then viewed under a widefield microscope (Leica DMIRB inverted microscope, USA). Fluorescent and phase contrast light microscope images were taken at x10 magnification. The Live/Dead Viability/Cytotoxicity assay was performed on cells cultured on the nanofibrous mats and control cells grown on standard tissue culture plastic. A negative control was also included; cells grown on plastic were incubated for 30 minutes with 70 % aqueous methanol to induce death and then the assay was performed.

MSC differentiation on composite cellulose:silk 75:25 nanofibres was determined through gene analysis. Nanofibrous discs were cultured with MSCs in stem cell expansion media supplemented with 5 or 10 ng/ml FGF-2. Tissue culture plastic controls were also included as follows; MSCs were cultured in standard 24-well tissue culture plates in stem cell expansion media supplemented with 5 ng/ml or 10 ng/ml FGF-2. MSCs were also cultured on plastic in chondrogenic differentiation media as a positive control for chondrogenesis. The chondrogenic differentiation medium consisted of high glucose DMEM containing 1 mM sodium pyruvate, 1 % ITS, 1 % P/S, 50 μg/ml ascorbic acid-2-phosphate and 1×10^−7^ M dexamethasone (all from Sigma, UK). This was supplemented with 10 ng/ml transforming growth factor-3 (TGF-β3, R&D Systems). A further control was also included – MSCs cultured in chondrogenic differentiation medium without TGF-β3. All cell cultures were incubated at 37 °C, 5 % CO_2_, 95 % air and media changed every 2-3 days. In all cases, cells were cultured for 14 days before downstream gene analysis.

RNA extraction was performed on composite fibres and tissue plastic control cell cultures after 2 weeks using the RNeasy Plus Mini Kit (Qiagen) and PureLink RNA Mini Kit (Thermo Fisher Scientific), respectively, following manufacturer′s instructions. RNA concentration and the purity of elutes was determined spectrometrically at 260 and 280 nm wavelengths (NanoPhotometer P-class Spectrophotometer, GeneFlow, UK). 20 ng/ml of RNA elution was reverse transcribed to produce complementary deoxyribonucleic acid (cDNA) using the High Capacity cDNA Reverse Transcription Kit (Applied Biosystems) according to the manufacturer’s protocol.

Chondrogenic gene expression was quantified using polymerase chain reaction (qPCR, StepOne Plus, Life Technologies). qPCR was performed on 96-well plates with each sample in triplicate. Each well contained 1 μl cDNA (of 20 ng/ml concentration), 3.5 μl nuclease-free water, 0.5 μl TaqMan^®^ Gene Assay primer and 5 μl TaqMan^®^ Gene Expression Master Mix. The following TaqMan^®^ Gene Expression Assays were used; Collagen type II alpha 1 (Col2), Aggrecan (Agg), SRY-box 9 (Sox-9), Collagen type I alpha 1 (Col1), Alkaline phosphatase (ALPL), Peroxisome Proliferator Activated Receptor Gamma (PPARG), and the housekeeping gene actin beta (ACTB) all purchased from Life Technologies Ltd. Data was analysed using the double delta Ct Analysis method. Gene expression levels were quantified relative to the expression levels of the housekeeping gene ACTB.

### 2.6. Statistical Analysis

Bone marrow MSCs were used from five patients in all experiments. For qPCR, each cDNA sample was run in triplicate, with the average being used for gene expression calculations. Statistical significance between gene expression levels of MSCs at different FGF-2 concentrations was assessed using the two-way ANOVA with Bonferroni post-test. *p*≤0.05 was taken as significant. Data points on graphs show mean ± standard error of the mean (SE).

## 3. Results and Discussion

### 3.1. Electrospinning Cellulose-Silk Composite Nanofibres

Cellulose and silk blend solutions were prepared in both TFA only and TFA-AcOH cosolvent systems at a range of concentrations and electrospun at various operational parameters. A 4 wt% concentration solution of the blended polymers in the cosolvent system electrospun at a 1.0 ml/hr flow rate and 2.0 kV/cm voltage was identified as ideal, generating a continuous electrohydrodynamic jet and resulting in the deposition of uniform nanofibres (*Figure 1A* *and 1B*). Further investigations revealed that the morphology of the nanofibres was affected by the environmental conditions, specifically relative humidity (*Supplementary Figure S1*). Environmental temperature and relative humidity (RH) were not controlled during the electrospinning experiments; however, they were recorded. The former parameter remained largely within a narrow range on all occasions, suggesting RH was playing the principal role in the significant differences in fibre diameter and morphology seen (*Supplementary Figure S2*). As RH increased, fibre diameters decreased, whilst bead length, width and frequency increased. Electrospinning occurs as a result of an electrically charged polymer jet being propelled in air, depositing as solidified fibres upon evaporation of its solvent medium. The impact of RH can be understood by considering the nature of both the solvent and polymer used in this process. As the electrospinning polymer jet emerges from the vertex of the Taylor cone, the charged jet briefly follows a straight flightpath until, due to bending instability, it enters a rapid whipping phase [34]–[36]. The latter phase enables the polymer jet to travel a length greater than the distance between the spinneret tip and grounded collector. As the jet spirals in the whipping phase, repulsive forces between charges carried by the jet cause elongation and thinning. Coupled with the evaporation of the solvent medium, this enables the deposition of fibres with diameters considerably smaller than the diameter of the spinneret that exuded the polymer solution. When using the highly volatile solvent TFA, an increase in RH is likely to cause rapid evaporation of this solvent. As the solvent evaporates, polymer chains begin to solidify, and intermolecular forces strengthen, overcoming jet elongational and thinning forces, resulting in larger fibre diameters. However, the relationship between RH and polymer chemistry must also be considered. Both cellulose and silk polymers are insoluble in water. An increase in RH reflects an increase in the amount of water vapour present in the air. Thus, the polymers precipitate faster, limiting elongational and thinning forces resulting in larger fibre diameters [37]. Furthermore, during electrospinning fibre beading can result due to increased jet instability in the whipping phase of the spinning process. Specifically, net charge density, surface tension and solution viscosity can influence the level of this instability [38]–[41]. As the polymer jet enters the whipping phase, it is not only exposed to elongational and thinning forces, but also to radial contraction forces [38], [39]. It is the latter force that causes fibre beading. The elongation and thinning forces on the jet must be sufficient to overcome radial contraction forces, otherwise beading will occur. If the polymer solution solvent evaporates rapidly during this phase, net charge density will be reduced, surface tension will increase and solution viscosity will decrease as the polymer chains solidify [39]. The results seen here are likely a combination of these competing factors at play. Environmental temperature remained within a narrow range in all instances (*Supplementary Figure S2)*.

**Figure 1.**
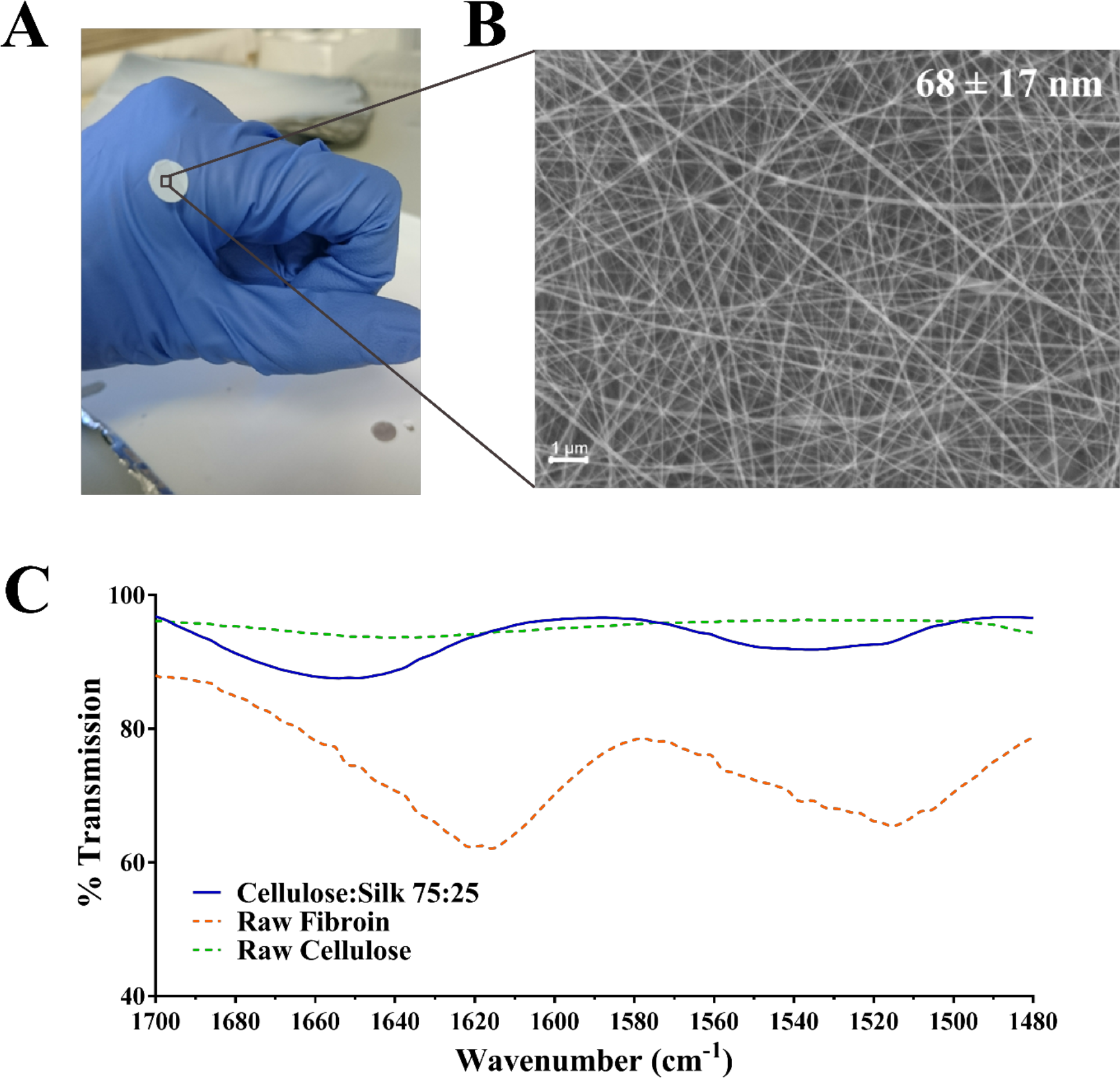
Electrospun Cellulose:Silk 75:25 Composite Nanofibres. Polymer solutions were electrospun at 1.0 ml/hr flow rate and 2.0 kV/cm voltage in a trifluoroacetic acid – acetic acid (TFA+AcOH) co-solvent system, A) 8 mm discs were cut for cell culture studies, B) Scanning electron microscopy of the nanofibrous material with fibre diameters of 68 ±17 nm (*scale bar inset measures 1 μm*), C) FTIR was performed to chemically characterise the nanofibrous mat. Solid line – electrospun fibres, dashed line – raw polymers. *Key shown inset*.

The impact of environmental conditions on the electrospinning process has not been widely investigated in the field of electrospinning research, however, it is beginning to receive attention [37] [42]–[45]. The influence of environmental temperature and RH on electrospun fibre morphology and diameter has shown dependence on solvent volatility and its miscibility with water and also polymer hydrophobicity [37], [43]. Although these experiments were not undertaken in a controlled environment, each parameter remained constant throughout. Strong trends were seen in resulting fibre morphology and diameter with reference to RH, whilst environmental temperature remained within a narrow range in all instances (*Supplementary Figures S1* *and S2*). This would need to be investigated in a controlled system before a firm conclusion can be drawn.

Chemical characterisation of the nanofibrous composite was performed using FTIR (*Figure 1C*). The regenerated polymers retained their native chemical functionalities. A strong broad band in the 3600-3000 cm^−1^ region denoting the O-H bond stretching of cellulose was present as was a narrower band at 2850-2920 cm^−1^, characteristic of asymmetric and symmetric stretching of methyl and methylene groups in cellulose, respectively [46]–[48]. The three characteristic regions of the silk fibroin peptide backbone are referred to as amide I (1700-1600 cm^−1^), amide II (1600-1500 cm^−1^) and amide III (1350-1200 cm^−1^) [49]–[51]. Strong absorption bands are seen in all three regions for the composite fibres (*Figure 1C*). The spectra of the composite polymer material showed broader bands at higher frequencies relative to the raw polymers. This can be attributed to changes in inter- and intramolecular hydrogen bonding, both within the protein and polysaccharide [46], [47], [51] as well as between the protein and polysaccharide [52]–[54]. Critically, the characteristic absorption bands of the solvents used – TFA and AcOH – were absent [26], [55].

### 3.2. Human Mesenchymal Stem Cell Viability on Composite Nanofibres

Nanofibrous mats with zero to minimal fibre beading were used in cell studies (≤ 20 beads per 100 μ^2^; *Supplementary Figure S2B*). In general, the presence of fibre beading in this range had no significant impact on cell binding, proliferation or differentiation. Human MSCs were cultured on the composite nanofibres with and without fibronectin (FN) coating. Fibronectin is an ECM glycoprotein involved in cell adhesion; enabling communication between intracellular and extracellular environments *via* cell surface integrin receptors [56], [57]. It is commonly used in biomaterial cell cultures to enhance cell adhesion to natural/synthetic material surfaces [32], [58]–[61]. The use of this adhesion protein does not conceal the surface functional groups of neat and composite cellulose-silk cast films, retaining their chemical influence on MSC behaviour [27].

Alamar blue (AB) assay was used to quantitatively assess hMSC viability and proliferation on the nanofibrous mats, with and without FN coating (*Figure 2A*). The AB assay was performed on the nanofibrous mats and tissue culture plastic control samples at three time points – day 1, 3 and 7 following initial cell culture (day 0). hMSCs were loaded at the same density on both surface types. Cells cultured on plastic surfaces remained viable and showed a gradual increase in cellular proliferation over the 7 day period. hMSCs cultured on nanofibrous mats maintained similar viability over the 7 day period, with and without FN coating (*Figure 2A*). This behaviour can be understood by considering the inherent chemical nature of the materials. Indeed, biomaterial surface chemistry can impact MSC adherence, differentiation and protein adsorption *in vitro*. The characteristic functional groups of the natural polymers used here - hydroxyl (-OH) for cellulose and amine (-NH_2_) for silk - have been widely studied, with their impact on MSC behaviour characterised [62]–[65]. Both −OH and −NH_2_ groups encourage MSC adhesion and proliferation through the promotion of cell surface integrins [64], with the latter group demonstrating a higher affinity [62]. Material surface chemistry can also influence the conformation of adsorbed proteins. Surface chemistry influences the conformation of adsorbed FN, and therefore it’s cell binding affinity, *via* the exposure of specific surface integrins [65]. Both −OH and −NH_2_ functionalised surfaces with adsorbed FN direct its conformation such that cell binding is enhanced. Furthermore, the strength of cell binding is greater on −OH functionalised surfaces [65].

**Figure 2.**
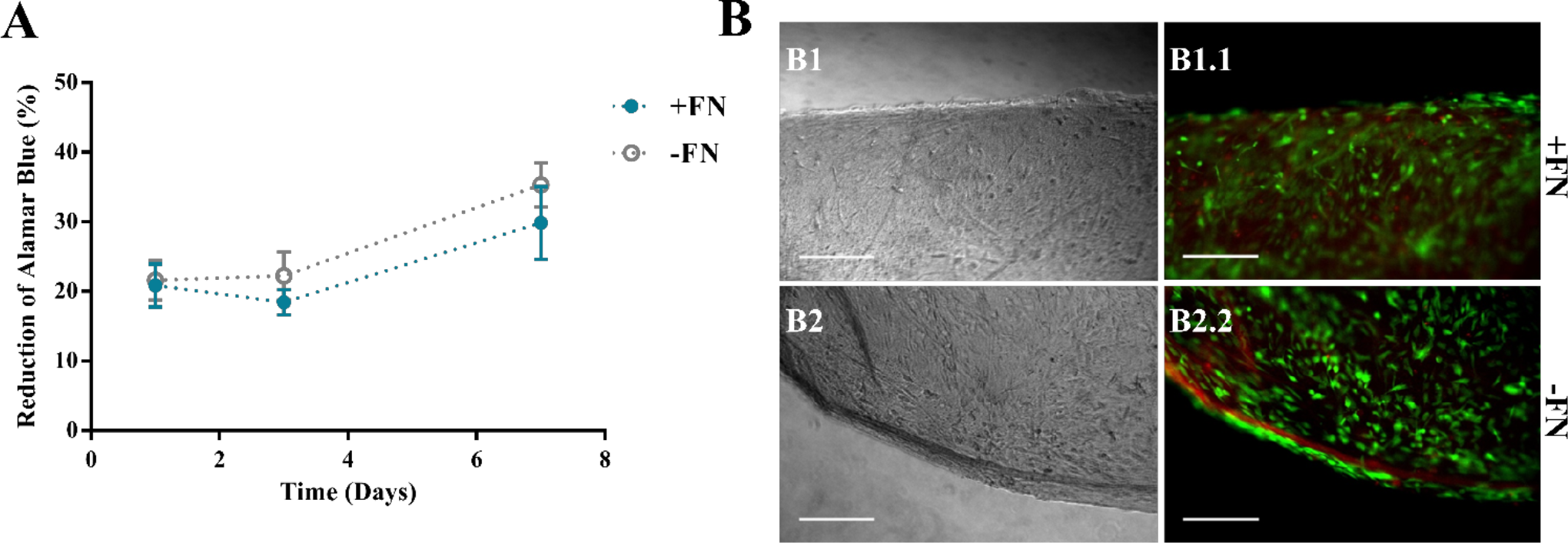
Viability of Human Mesenchymal Stem Cells on Composite Nanofibres. Cells were cultured on nanofibrous discs with (blue line) and without (grey line) fibronectin (FN) coating and their viability assessed. A) Viability at 1, 3 and 7 days using Alamar Blue assay, data points show mean ± SE (n=5). B) a LIVE/DEAD viability assay was performed at day 7. B) Optical (B1 and B2) and fluorescence (B1.1 and B2.2) microscopy images of cells on composite nanofibres. Live and dead cells show green and red fluorescence, respectively. Scale bar inset measures 300 nm. Representative images shown (n=5).

Although the use of this ECM glycoprotein encouraged MSC adhesion to these surfaces, there was statistically no significant difference between MSC adhesion on nanofibres without this coating (*Figure 2A*). It can therefore be concluded that use of this adhesive protein does not mask the surface functional groups of these natural polymers, and therefore their respective roles in cell binding affinity. The TFA-AcOH cosolvent system used in their fabrication did not elicit any detrimental effects, as anticipated from the chemical characterisation performed on the materials (*Figure 1C*). Regardless of FN coating on the nanofibres, the reduction of AB (and therefore cellular activity) increased over the 7 day period, demonstrating not only the biocompatibility of these materials but also their ability to support cellular proliferation.

The non-toxic nature of the AB assay enabled further qualitative analysis to be performed on the same samples to visualise cell adherence on nanofibrous and plastic surfaces. Viable cells were viewed on all surfaces, corresponding to the AB assay results at day 7 (*Figure 2A*). The MSCs grown on tissue culture plastic, as a positive control, demonstrated a spindle morphology, characteristic of healthy undifferentiated MSCs (*Supplementary Figure S3*). The presence of dead cells was negligible. MSCs killed with methanol, used as a negative control, all showed red fluorescence demonstrating their compromised membrane integrity, with loss of their spread spindle morphology (*Supplementary Figure S3*). For MSCs grown on the composite nanofibres, images were taken at the scaffold edge and centre to visualise whether cell spread was uniform across the 8 mm disc. In all cases, cells were evenly spread across the nanofibrous surface, regardless of FN coating. Cell morphology was more compact than cells grown on plastic with negligible dead cells (*Figure 2B*).

To visualise cell morphology at a higher resolution on the composite nanofibres, following 7 days of *in vitro* culture the cell-laden biomaterials were fixed and viewed under SEM (*Figure 3*). Densely populated cell layers with clear cell-to-cell contact were seen on the materials, regardless of FN coating. Critically, the fibres had retained their nanofibrous morphology and the mats retained their overall shape and integrity until the end of the *in vitro* culture.

**Figure 3.**
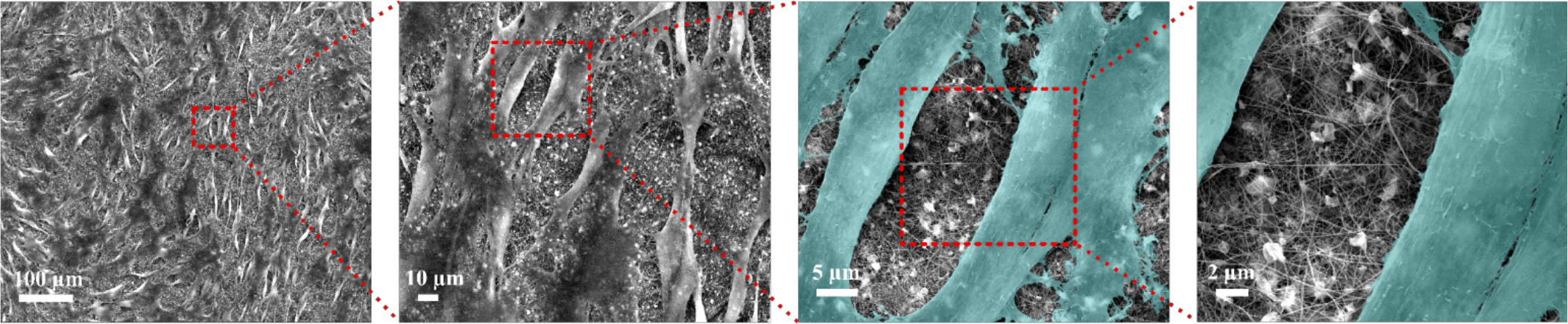
Mesenchymal Stem Cells on Composite Electrospun Mats. Human stem cells were cultured on cellulose-silk composite nanofibres for 7 days. Samples were then fixed and viewed using scanning electron microscopy. Representative image shown with increasing magnification, *n = 5. Cells were coloured in latter images to enhance contrast between cells and nanofibres*.

### 3.3. Human Mesenchymal Stem Cell Chondroinduction on Composite Nanofibres

Having established hMSC viability and proliferation on these nanofibrous materials, and therefore their biocompatibility, the stimulative capacity of the scaffold was investigated, as cast films of cellulose and silk blended in a 75:25 mass ratio have been shown to be chondroinductive [27]. To this avail, hMSCs were cultured on FN-coated nanofibrous composites in the absence of stimulating factors. Cells were also grown on tissue culture plastic under the same media conditions as control. Cells were cultured in 5 ng/ml and 10 ng/ml FGF-2. FGF-2 is used in hMSC culture *in vitro* due to its ability to enhance cellular proliferation and maintain stem cell differentiation potential [28], [29] . It is not known to behave as a chondrogenic stimulator. As a further control, hMSCs were cultured on the plastic surface in the presence of transforming growth factor-ß (TGF-ß) (*Supplementary Figure S4*). This soluble growth factor is a potent stimulator of chondrogenesis in MSCs and is the gold standard for this purpose *in vitro* [66]. To confirm the specific chondrogenic differentiation of hMSCs on the composite biomaterial, gene expression for other potential lineages was measured. This included the osteogenic marker (ALPL) and the adipogenic marker (PPARG) (*Supplementary Figure S5*).

Human MSCs on FN-coated nanofibrous and plastic surfaces maintained viability over the longer investigation period (14 days). Following the culture period, downstream analysis was performed to screen for the expression of chondrogenic genes on all surfaces (*Figure 4*). The expression levels of type-II collagen (Col-2), aggrecan (Agg) and SRY-box 9 (Sox-9) were assessed, as key markers for chondrogenesis in hMSCs [67]–[70]. Type-I collagen levels were also assessed as a marker unusually associated with hMSC chondrogenic differentiation *in vitro* [71]–[73]. Expression levels of all genes were normalised to that of the housekeeping gene ß-actin [74].

All the chondrogenic genes studied on the nanofibrous composites were upregulated, relative to the hMSCs control grown on plastic (*Figure 4*). This upregulation was statistically significant at the lower FGF-2 concentration of 5 ng/ml. These results strongly indicate that the nanofibrous composite surface can influence hMSC behaviour and direct its differentiation without the need for stimulating factors. Importantly, the significance of gene upregulation is influenced by the concentration of FGF-2. The latter is likely due to the hMSCs being under the influence of a higher proliferative stimulus, thereby overriding hMSC capacity to differentiate. At the lower FGF-2 concentration however, the inherent material characteristic driving hMSC differentiation appears to overtake FGF-2 proliferative effect. Taken together, these results demonstrate the potent capacity of the nanofibrous cellulose/silk composites to drive chondrogenic differentiation independent of soluble chondrogenic growth factors, making these composites exciting candidates for tissue engineering applications. Here, the introduction of a biomimetic nanofibrous matrix resulted in a more potent impact on chondroinduction than that previously reported on cast films of the same composition [27].

**Figure 4.**
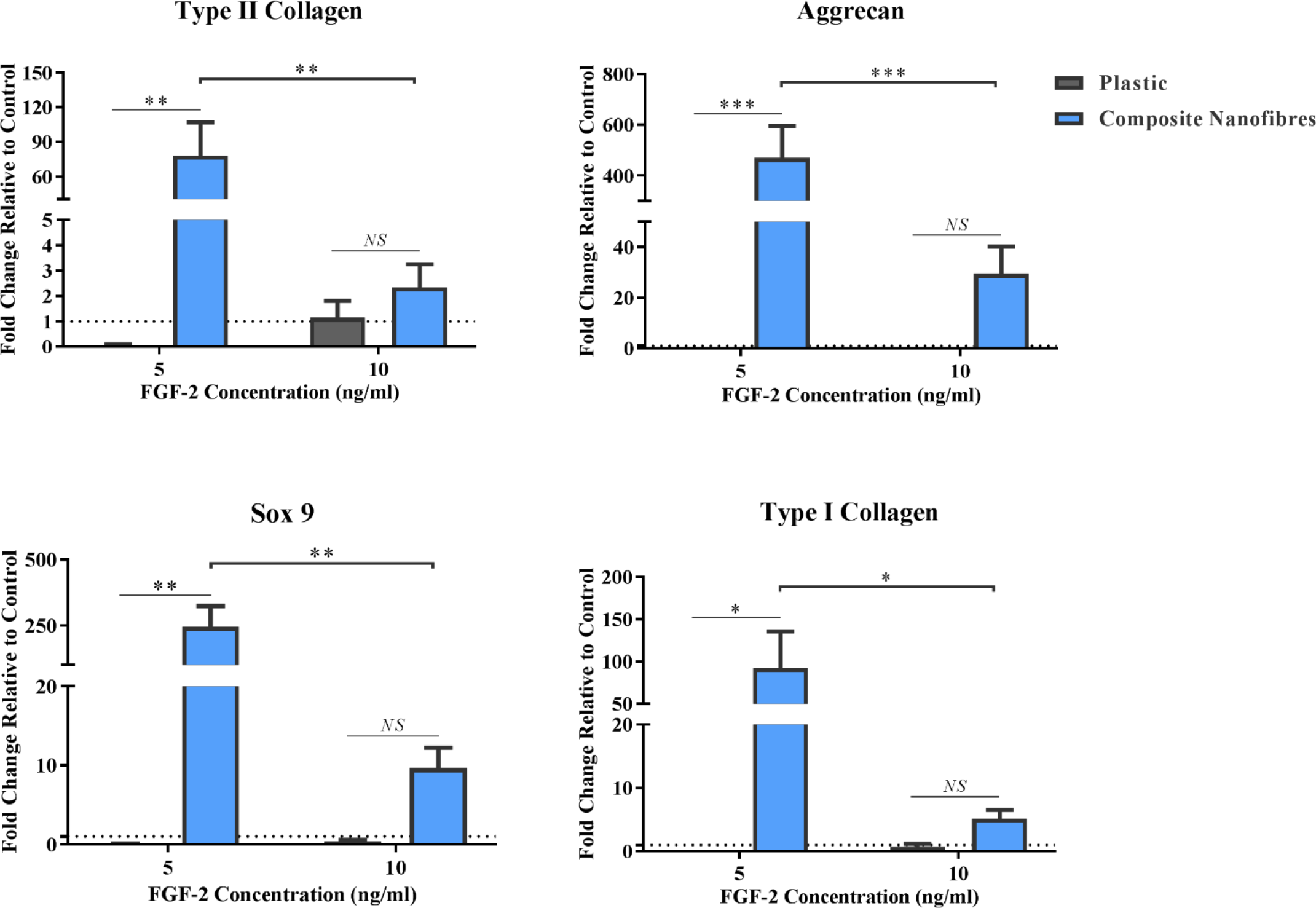
Stimulative Capacity of Composite Nanofibrous Mats. Human mesenchymal stem cells (MSCs) were cultured on plastic and composite nanofibres in the absence of soluble chondrogenic factors. Following 14 days culture, qPCR was performed to quantify the level of chondrogenic genes expression - Col2, Agg and Sox-9 - as well as Col1. Gene expression levels were normalised to the expression of the housekeeping gene, ACTB (shown in dotted line). Graph shows mean ± SE, *n*=5. *Samples were compared statistically using two-way ANOVA*. * = *p<0.05*, **=*p<0.001*, ***=*p0.0001*.

It is important that the specific inherent material characteristic implicated is understood. This will not only assist future biomaterial design considerations, but also introduce the possibility to control and tune the level of gene expression induced. Recently, investigations were carried out on cast films of these natural polymer composites cultured with hMSCs [75]. The chondroinduction of hMSCs was significantly reduced using blebbistatin – a potent inhibitor of cell force transmission between intra- and extra-cellular environments. This suggests that the mechanical rather than the chemical properties of the material are the key drivers of its chondroinductive behaviour.

The impact of electrospun fibre diameter on cell growth and differentiation *in vitro* is well-established [4], [76]–[81]. Electrospun fibres of nano- and micro-scale appear to support chondrogenic differentiation (in the presence of stimulating factors) in a context dependant manner [4], [76], [77], [80], [81]. Human MSCs cultured in the presence of stimulating factors show a preference for micro-scale fibres when undergoing chondrogenesis [4], [76], whereas nano-scale fibres are favoured by chondrocytes – the terminally differentiated cells exclusive to cartilage tissue [80], [81]. Therefore, electrospinning the composite cellulose:silk 75:25 blend to fabricate microfibers for *in vitro* culture with hMSCs may enhance the composite chondroinductive capacity proposing an additional potential mechanism for tuning the materials smart behaviour. Chondrocytes undergo dedifferentiation over long-term *in vitro* culture, limiting their use in cartilage tissue engineering due to low cell yields [82]–[84]. Perhaps culturing these cells on a potentially chondroinductive nano-scale smart material will help maintain their chondrogenic phenotype, without the need for growth factor supplements – enabling their long-term culture *in vitro*.

## 4. Conclusion

Cellulose and silk composite nanofibres have been directly electrospun in a 75:25 mass ratio for the first time, with no post-spun treatments required. The morphology of these novel nanomaterials could be tuned through adjustments in operational and environmental parameters. Their biocompatibility has been demonstrated using hMSCs, supporting not only cell proliferation but also the ability to drive chondrogenic differentiation, without the need for stimulating factors. It is important to highlight however that the electrospun biomaterials clearly lack sufficient porosity to aid cell infiltration (*Figure 3*). Indeed, this is a common challenge when employing an electrospinning fibre fabrication technique. Ensuring porosity is sufficient to support cell infiltration and spread is key to developing an adequate material for tissue engineering applications. Thus, future design considerations would need to address this limitation.

These findings are significant not least due to the use of these abundant natural polymers; this ratio directed hMSC differentiation down a specific lineage in the absence of stimulating factors. Currently, research in the field of cartilage tissue engineering using biomaterials requires the use of stimulating factors to drive the chondrogenic differentiation of MSCs on these surfaces [85]–[89]. Such stimulants are often costly, have limited efficiency, are short-lived, lead to terminal differentiation and are animal derived. These limitations beg the case for smart biomaterials with an inherent instructive capacity.

We have demonstrated the successful reproduction of a chondroinductive natural polymer composite into a more biomimetic configuration. Using the native ECM architecture to guide our biomaterial design, we have fabricated biocompatible nanofibrous networks with an inherent capacity to drive stem cell differentiation. Future work will focus on enhancing the 3D component of these materials; to aid cell infiltration and enable mechanical support for the engineered tissue. Current findings coupled with our studies investigating the biochemical signalling pathways implicated in driving this behaviour [75] will help inform future smart biomaterial design strategies; bringing us closer to addressing the growing need for adequate treatments for degenerative joint diseases.

## Acknowledgments

The authors would like to thank the Engineering and Physical Sciences Research Council (EPSRC) for funding this work through a Doctoral Training Partnership grant EP/K502996/1 and a Doctoral Prize grant EP/N509619/1. RB would also like to thank Professor Mark Lowenberg (Postgraduate Tutor) for the support provided during the Doctoral Training Partnership.

## Author Contributions

R.B. designed and performed the experiments and wrote the manuscript, B.S. supported the electrospinning work, A.W.P. and F.S. contributed to the experimental design. W.K. conceived the original idea, supervised the project and wrote the manuscript. All authors provided critical feedback and helped shape the research, analysis and manuscript.

## Supplementary Info

**S1.**
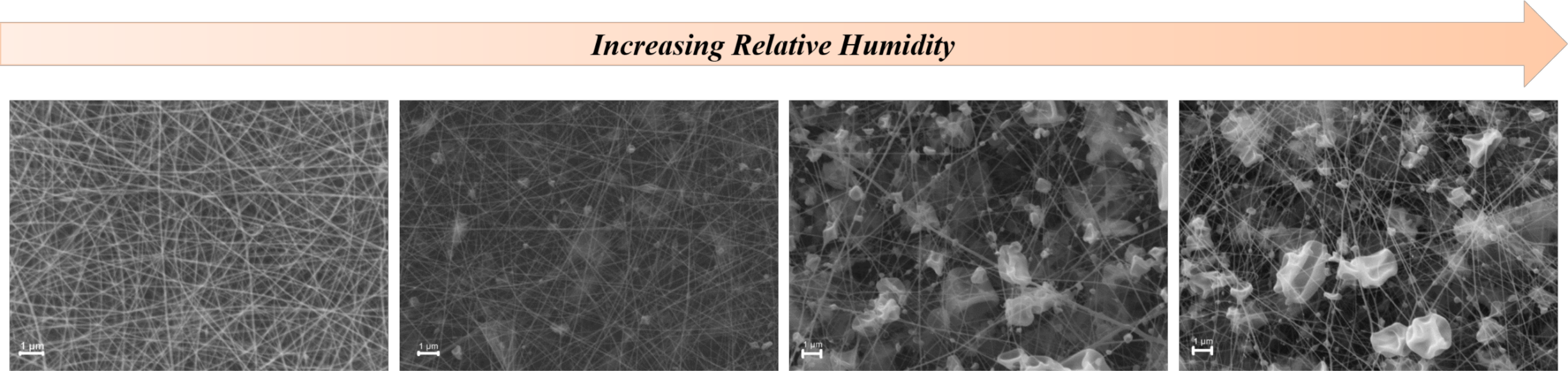
Electrospun Cellulose:Silk 75:25 Composite Nanofibres. Polymer solutions were electrospun at 1.0 ml/hr flow rate and 2.0 kV/cm voltage in a trifluoroacetic acid - acetic acid (TFA+AcOH) co-solvent system. An increase in environmental relative humidity affected fibre morphology. Scale bar inset measures 1 μm

**S2.**
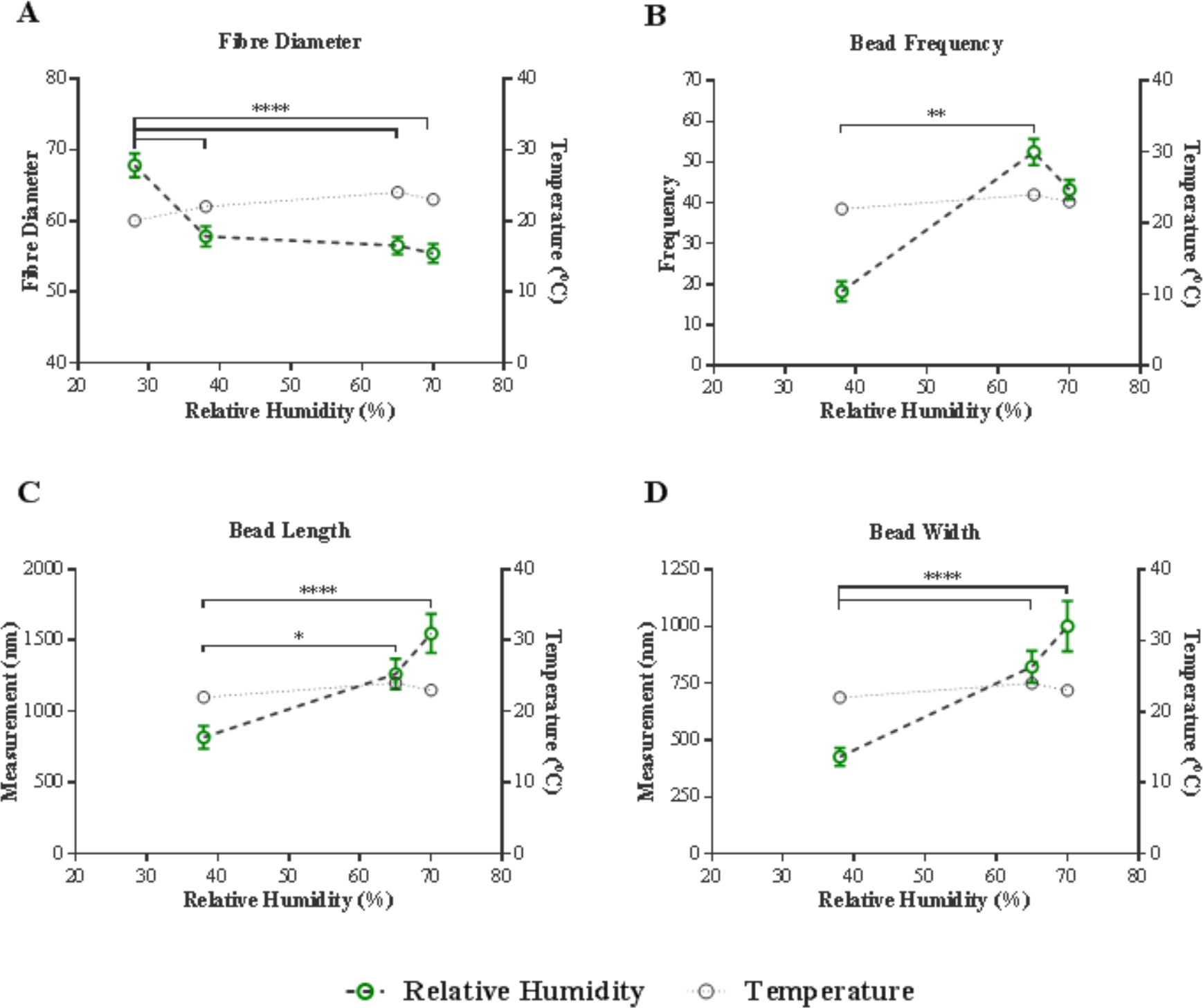
Cellulose:Silk 75:25 Composite Nanofibres. Polymer solutions were electrospun at 1.0 ml/hr flow rate and 2.0 kV/cm voltage in a trifluoroacetic acid . acetic acid (TFA-AcOH) cosolvent system. Graphs show impact of environmental humidity on A) Fibre Diameter, B) Bead Frequency, C) Bead Length and D) Bead Width. Fibre analysis shows mean of 100 measured fibres. Bead frequency analysis shows mean of four separate areas, each measuring 100 μ^2^. Bead analysis shows mean of 20 measured beads. All graphs show Mean ±SE (Left y axis, green). Non-parametric Kruskal-Wallis test with Dunn′s test post hoc applied to all graphs. *p*≤0.05 taken as significant. Environmental temperature remained within a narrow range across all samples (Right y axis, grey).

**S3.**
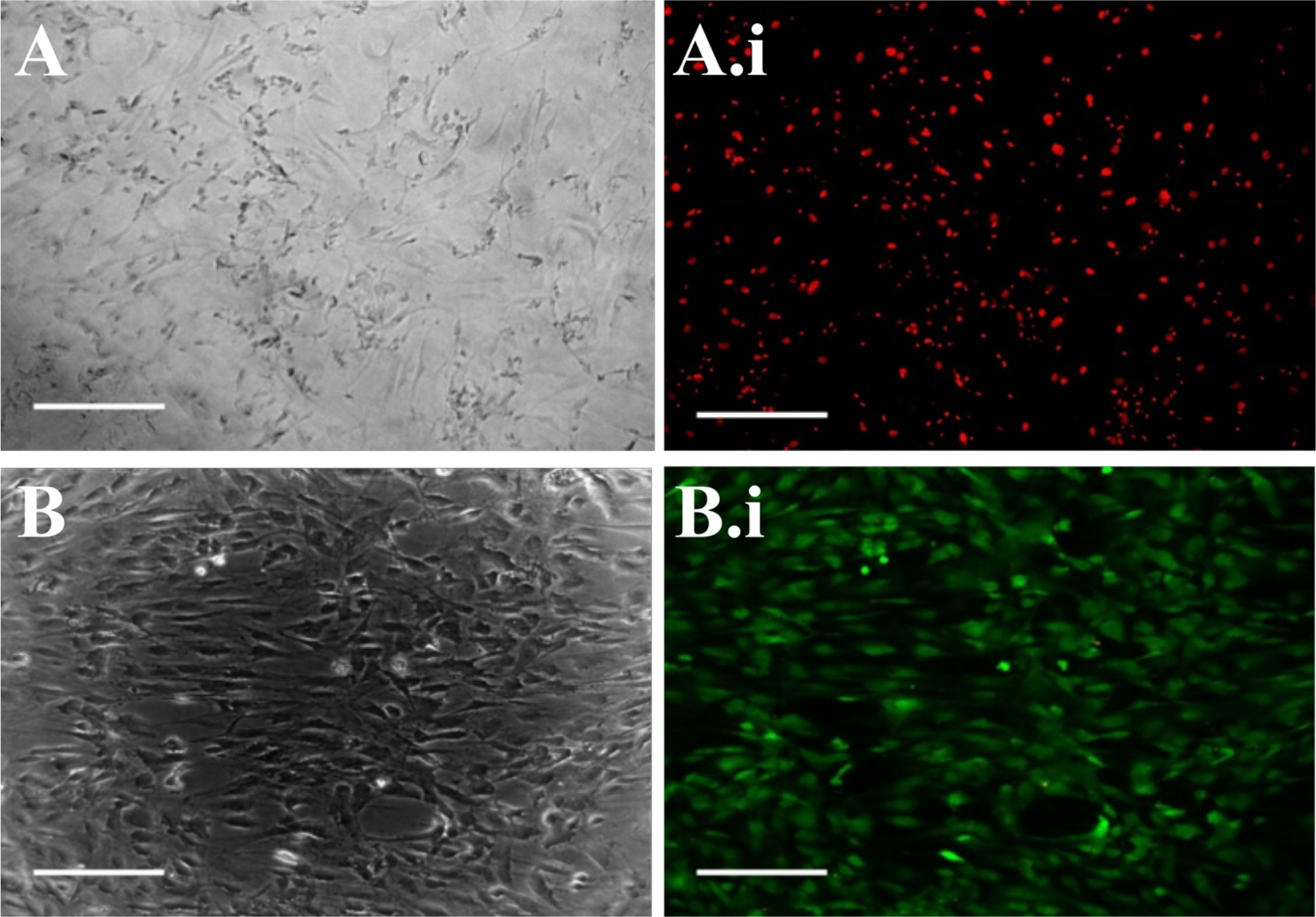
Viability of Human Mesenchymal Stem Cells on Plastic. Cells were grown on tissue culture plastic and their viability assessed after 7 Days using a LIVE/DEAD viability assay. A.i), B.i) Optical microscopy and A.i), B.i) Fluorescence microscopy images of cells. Images were taken at x10 magnification. Scale bar inset measures 300 nm. Representative images shown of cells from five patients. Live and dead cells show green and red fluorescence respectively.

**S 4.**
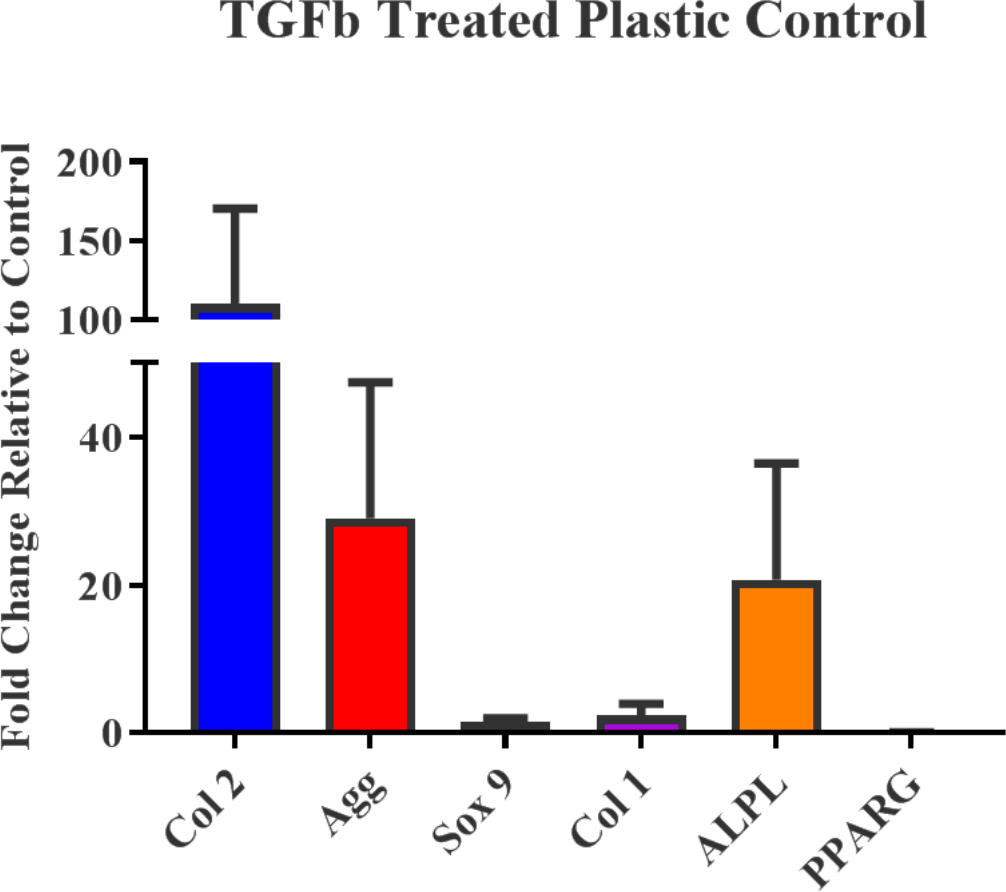
Human Mesenchymal Chondroinduction. Human stem cells were cultured on plastic in the presence of chondroinductive media. Cells were screened for the expression of key chondrogenic (Col 2, Agg, Sox 9, Col 1), osteogenic (ALPL) and adipogenic (PPARG) genes 21 days later. Gene expression levels were normalised to the expression of the housekeeping gene, ACTB (shown in dotted line). Graph shows mean ± SE, *n*=5.

**S 5.**
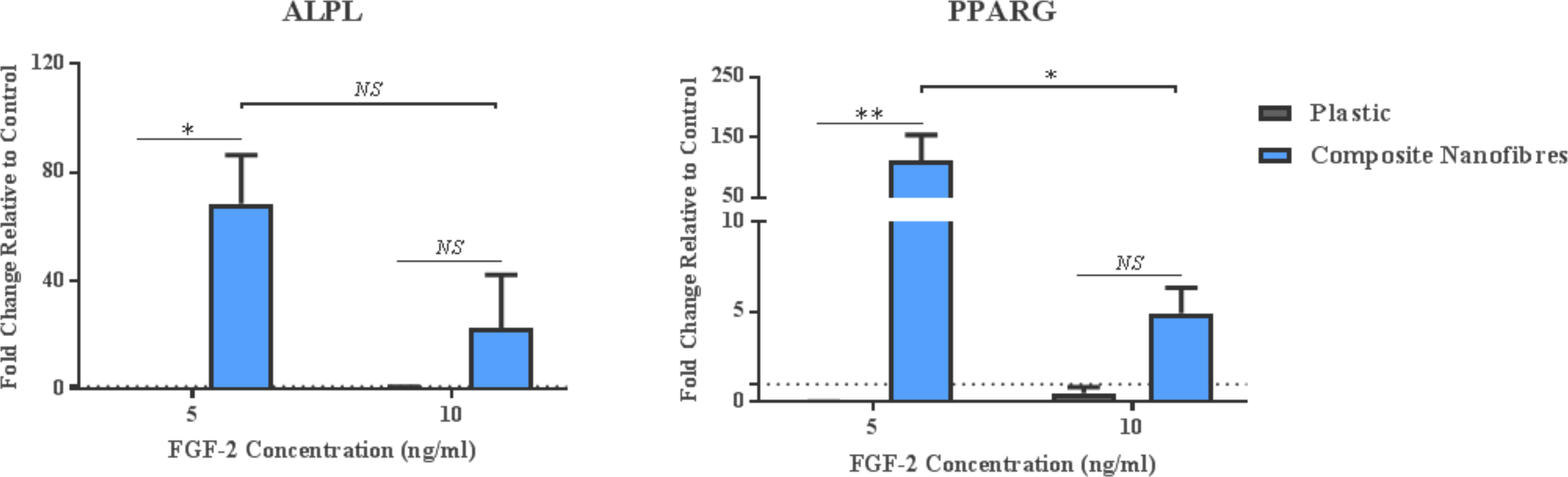
Stimulative Capacity of Composite Nanofibres. Human mesenchymal stem cells were cultured on plastic and composite cellulose-silk nanofibrous mats in standard mesenchymal stem cell expansion media supplemented with fibroblast growth factor −2 (FGF-2) at two concentrations. Cells were screened for the expression of key osteogenic (ALPL) and adipogenic (PPARG) genes 21 days later. Gene expression levels were normalised to the expression of the housekeeping gene, ACTB (shown in dotted line). Graph shows mean ± SE, *n*=5.

